# New technologies for hair follicle growth and regeneration: comparison between different commercial products contain exosomes

**DOI:** 10.1101/2024.11.17.623990

**Authors:** Greta Ferruggia, Massimo Zimbone, Maria Violetta Brundo

## Abstract

In people suffering from baldness, stem cells remain inactive and are unable to regenerate new hair. For this reason, research has focused on factors (including growth factors and cytokines) or new technologies that could favor the transition from telogen to anagen and the subsequent stimulation of hair growth and regeneration. In our study we compared *in vitro*, on cultures of human hair follicles, the effect on hair growth and on the regeneration of the dermal papilla of two products containing exosomes of different origins obtained with innovative technologies. In particular, we used a product in which the presence of exosomes and proteins obtained from stem cells derived from the umbilical cord lining is declared and a product that contains exosomes obtained from bovine colostrum and passively loaded with a mixture of growth factors and cytokines purified from bovine colostrum. The analyzes demonstrated a notable increase in the growth and regeneration of the bulbs the dermal papilla in the samples treated with the product containing exosomes obtained from bovine colostrum compared to product containing exosomes from mesenchymal stem cells which was comparable to the result obtained in the control samples (untreated). Therefore, the use of products containing exosomes, growth factors and cytokines purified from colostrum could be successfully used as possible new treatments capable of significantly slow hair loss and promote the growth of new hair.

## 1. Introduction

Hair-care products occupy an important market share in the cosmetic sector worldwide, a direct consequence of the fact that, nowadays, hair plays a decisive role in the life of every woman or man [1,2]. In a market that is always attentive to new consumer needs, often governed by aggressive marketing, it has today become a very important prerogative in providing a product that is as green as possible, both towards the environment and the person who uses it, but at the same time that it does not forget to perform its function as best as possible, the cardinal point of every self-respecting product [1-3]. The market for cosmetic raw materials for hair care is in fact developing towards products which certainly perform well on the hair fiber, but which at the same time respect and preserve the skin of the scalp, or even which also provide beneficial effects for the scalp, always within the limits of the cosmetic product. The cosmetic is not a drug: it cannot boast a therapeutic effect on the skin but it can certainly improve and preserve its appearance, help restore its structure and rebalance its functionality [4].

Hair growth is cyclical, made up of a growth phase, Anagen, a fall phase, Catagen, and a rest period, Telogen. Anagen is the growth phase, i.e. the beginning of the mitotic activity of the cells of the follicle matrix which can last from two to seven years [5]. It is divided into 6 sub-phases. The first 5 employment for a relatively short period and constitute the proliferative phase. The sixth period is certainly the longest as it represents the differentiation phase with different times in men compared to women. Approximately 85-90% of all hair is in this phase. Catagen is the cessation of metabolic activity, therefore of growth; it lasts about two weeks during which the keratinized bulb remains linked to the papilla through a sac formed by the external epithelial sheath which contains the last cells produced in the mitotic phase. The end of this phase is given by the detachment of the papilla from the sac. It is estimated that 1% of total hair is in this phase. Telogen lasts five to six weeks during which the hair does not grow and remains attached to the follicle. When the hair falls out the growth cycle begins again. The percentage of hair in Telogen is 10-20% [5]. There are mechanisms for regulating the growth cycle and its phases: one is hormonal control, which is the main cause of androgenic alopecia, the second is the control of growth factors [6]. Messenger and Collaborators [6] have clarified that as regards growth factors, the different types of Epidermal growth factor (EGF) act on the external follicular sheath and some of them, such as hair growth factor (HrGF) or fibroblastic growth factor 5 (FGF 5), have a stimulating effect on hair growth [6].

Alopecia is defined as hair loss anywhere on the body [3]. Alopecia can be divided into cicatricial alopecia and non-scarring alopecia. Cicatricial alopecia is linked to the active destruction of the hair follicle, which, when irreparably damaged, is replaced by fibrous tissue. This form of alopecia sees permanent hair loss and there are two subtypes: primary and secondary. Instead, non-scarring alopecia is due to processes that slow or reduce hair growth without creating irreparable damage to the hair follicle. Among the different types of non-scarring alopecia, there are telogen effluvium, alopecia areata, anagen effluvium and androgenetic alopecia. Basic research in hair follicle biology is defining potential new avenues for treatment of hair loss conditions and methods to induce new hair growth [7]. New therapies involving biomolecules and exosomes are also among the emerging possibilities for treatment of alopecia [8]. Aims of the present study was to compare *in vitro* effects of some commercial compounds on hair follicles growth and regeneration. We have tested the effectiveness of products contain exosomes from human stem cells of umbilical cord lining (CSCs-Exo) and products contain bovine colostrum exosomes loaded with growth factors and cytokines purified from bovine colostrum (Colostrum-Exo). Stem cell exosomes have already been extensively studied and characterized by several Authors [8-10], but however, there are still some limitations in this field that need to be attended [8]. In fact, some Authors suggest further studies are needed to investigate the strategies associated with exosomes to improve their clinical efficacy in the field of hair growth [8]. Despite favorable outcomes from many of the preclinical studies, however, no clinical studies were reported employing exosomes from CSCs for hair growth to be due to the high production cost and requirement of specialized purification methods associated with their short shelf life [11]. In addition, the administration of stem cell-derived exosomes carries the potential risks of the uncontrolled transmission of genetic information and immune responses [11]. Conversely, from bovine milk or bovine colostrum can generate massive amounts of exosomes cost-effectively compared to exosomes derived from cell lines [12,13]. These exosomes are highly appreciated for therapeutic applications in terms of safety [11]. In addition, colostrum exosomes have been found to contain over 1,000 growth factors and biological signals such as FGF, PDGF, VEGF and cytokines that were critical for promoting and accelerating healing, tissue regeneration [14,15]. Exosomes from bovine colostrum thus are considered as a new effective non-surgical therapy for hair loss. Results of several studies showed that colostrum exosomes promoted the proliferation of Human hair dermal papillary cells and rescued dihydrotestosterone (DHT, androgen hormones)-induced arrest of follicle development [11]. The Authors showed that colostrum derived exosomes accelerated the hair cycle transition from telogen to anagen phase by activating the Wnt/β-catenin pathway [11].

## 2. Materials and Methods

For our research we purchased commercial products used for hair follicles growth and regeneration. In particular, one product contains exosomes from mesenchymal stem cells extracted from the lining of the umbilical cord, fibronectin, glycoproteins, albumin, collagen, hyaluronic acid (indicated with CSCs-Exo). The second product contains exosomes purified from colostrum and passively loaded with growth factors and cytokines derived purified from colostrum together with hyaluronic acid, *Curcuma longa, Callus* extract, Vitamins, amino acids, peptides and natural elements (Mg, Zn, Na, Mn) (indicated with Colostrum-Exo).

### 2.1. Human hair follicle *in vitro* culture

Human follicles from of 10 healthy male individuals were micro-dissected and isolated from the occipital scalp during hair transplantation procedures. All individuals signed informed consent, according to the Helsinki Declaration (2001). Follicles in anagen phase VI were selected via stereomicroscopy and were cultured in 24-well plates (1 per well) with 500 μL Williams medium (Gibco BRL, Rockville, MD, USA), with insulin 10 μg/mL, hydrocortisone 10 ng/mL, streptomycin 10 μg/mL, and penicillin 100 U/mL (Wuppertal, Germany), at 37 °C and 5% CO_2_. A total of 72 follicles were distributed in different plates to test the two products previously dissolved in Williams medium. The growth parameters of the follicles were analyzed every 3 days for 18 days. Hair length was assessed using an inverted microscope (LEICA DM IRB), and the images obtained were measured using ImageJ. Three replicates were performed.

### 2.2. Identification and analysis of hair follicle derma papilla cells

After 18 days of exposure, hair follicles were stained with DAPI (Sigma-Aldrich) to highlight the nuclei in blue and allowed them to be identified. Hair follicles were immediately visualized under the fluorescence microscope (Nikon Eclipse Ci, Melville, NY, USA).

### 2.3 Isolation and characterization of exosomes

Exosomes present in the two products were analyzed by a light-scattering system according to [17,18]. Measurements were performed using a quartz scattering cell, confocal collecting optics, a Hamamatsu photomultiplier mounted on a rotating arm, and a BI-9100AT hardware correlator (Brookhaven Instruments Corporation, Holtsville, NY, USA). The samples were illuminated with a 660 nm laser. A low power intensity was used (5-15 mW) to avoid convective motions due to local heating.

### 2.4. Statistical analysis

The analyzed parameters were represented by mean ± standard deviation. Statistical and multiple comparisons of the data were performed by ANOVA, followed by Tukey’s test, considering p values less than 0.05 as significant. Significant data were represented with the symbol * (p < 0.01), while strong statistical differences were reported with the symbol ** (p < 0.001).

## 3. Results and Discussion

Dynamic light scattering (DLS) is an optical analysis method for measuring the size and distribution of submicron particles [18]. Light scattering measurements were performed by a homemade apparatus using a low power intensity was used to avoid convective motions due to local heating, because of the Brownian motion of the particles in solution, the light scattered by particles suspended in solution fluctuates in time. The intensity auto-correlation function g2 provided by the hardware correlator operating in “single photon counting regime” is a decreasing function with a well-defined baseline. The field autocorrelation function g1 is, thus, calculated by the g2 using the Siegert relation g2=|g1|^2^. For polydisperse noninteracting particles in Brownian motion, the g1 shows several exponential decay components which are analyzed by the cumulants method, thus, following the standard theory of DLS, it is possible to calculate the hydrodynamic diameter [17,18]. Light scattering analysis showed that in the product that should contain mesenchymal stem cell exosomes (CSCs-Exo) there are nanometric structures, but the detected structures have dimensions of 687 ± 1.8 nm, so with a range greater than 40 and 160 nm (on average 100 nm) in diameter, typical dimensions of exosomes. The concentration of these structures was not detectable by the instrument (Fig.1A). Instead, exosomes were obtained from product with bovine colostrum (Colostrum-Exo) have the average diameter size was 112 ± 1.2 nm and presented a spherical morphology. The concentration mean was 3.8 × 10^9^ particles/mL (Fig.1B).

**Figure 1.**
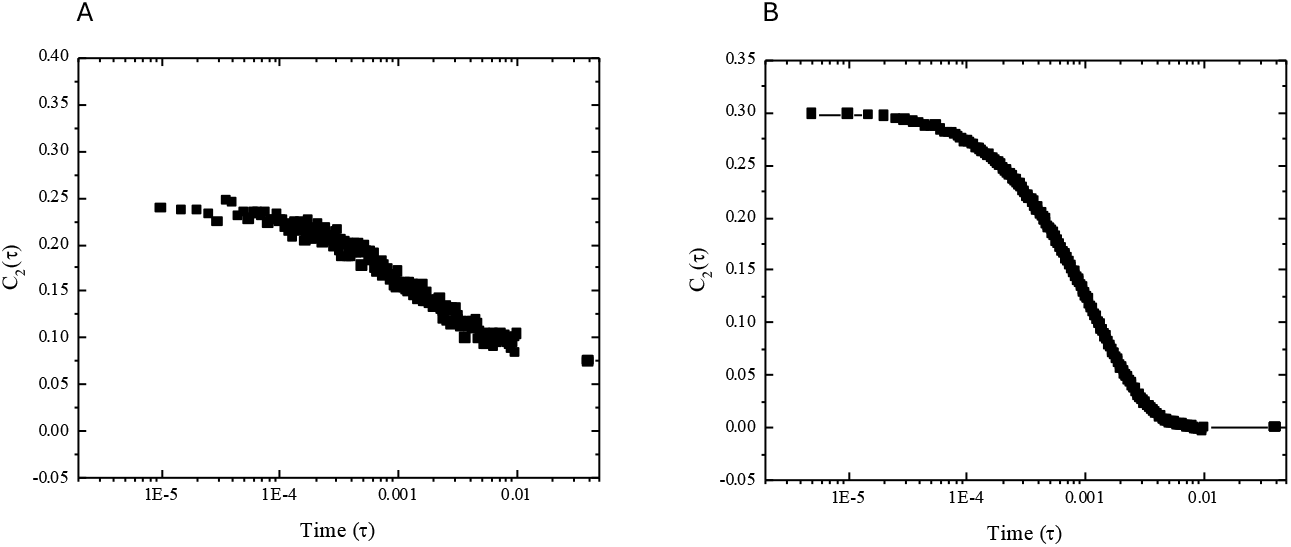
DLS analysis was used to characterize the size distributions of exosomes within two products: (**A**) product with exosomes from mesenchymal stem cells and (**B**) product with exosomes from colostrum bovine loaded with growth factors and cytokines purified from bovine colostrum.

The test used for to verify the growth of the hair follicle in the presence or absence of the products to be tested showed samples exposed to product having CSCs-Exo don’t have a significant difference in the width of germinative growth area of the hair bulb compared to the negative control (Fig. 2). The samples treated with product containing Colostrum-Exo, showed higher growth (Fig. 2). Hair follicles growth was done for 18 day and their length measured at regular every 3 days. The results showed the effectiveness of product contains exosomes purified from colostrum and passively loaded with growth factors and cytokines derived from colostrum compared with product containing exosomes from mesenchymal stem cells (Figure 2). Specifically, at the end of treatment, follicles exposed to the product containing Colostrum-Exo showed a length of 3,19 ± 0.12 mm. Lower growth was shown following treatment with CSCs-Exo, where the length corresponded to 2.11 ± 0.9 mm. Finally, the negative control showed a value of 1.73 ± 0.8 mm. The treatment with product indicated with Colostrum-Exo showed a statistically significant difference (p < 0.01) in the first 12 days and highly significant in the checks carried out on the 15th and 18th day (p < 0.001), compared with the untreated samples (negative control) and samples treated with product containing exosomes from mesenchymal stem cells (Figure 2).

**Figure 2.**
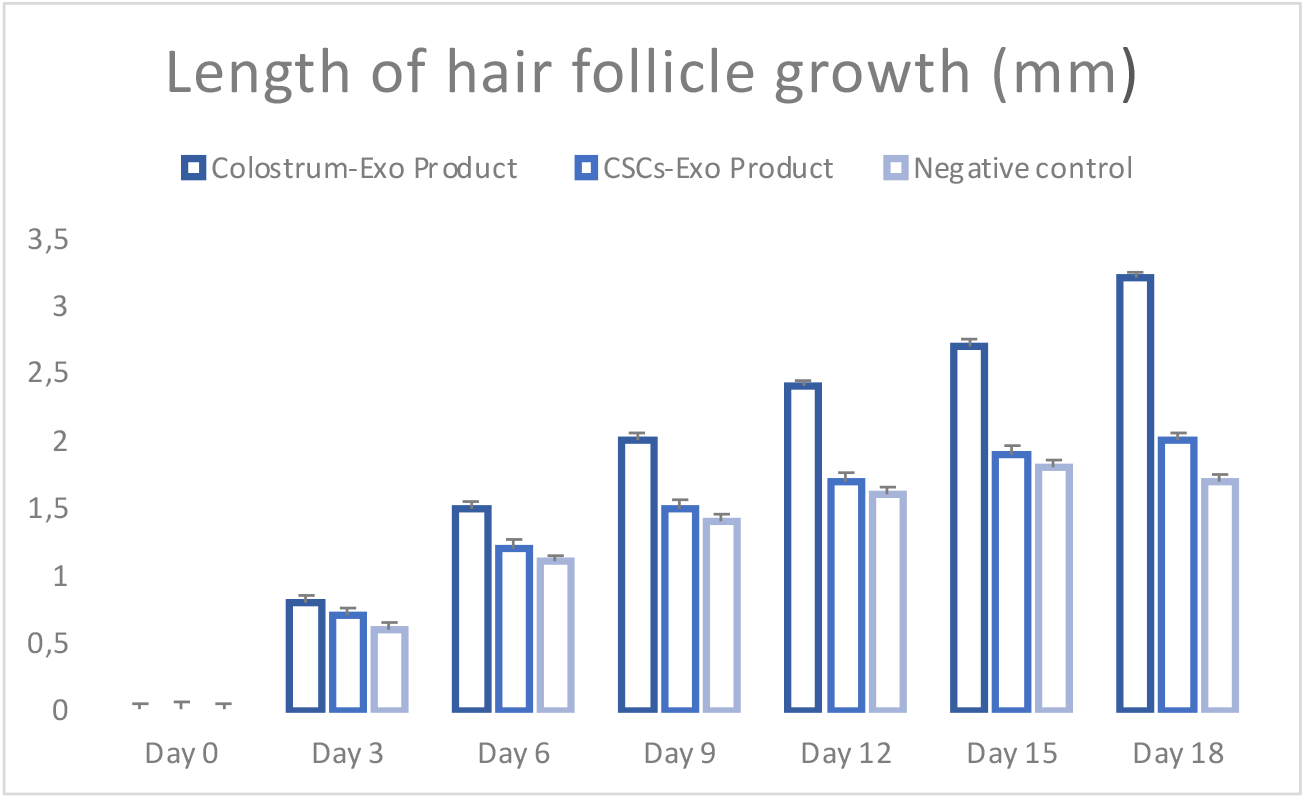
Comparison of hair length variation of samples exposed to different products. Statistical differences were indicated with the symbol * (p < 0.01) or ** (p < 0.001) for more significant.

The expansion of the germinative area in bulbs treated with product containing exosomes from colostrum bovine loaded with growth factors and cytokines purified from bovine colostrum was greater compared to samples treated with product having exosomes from mesenchymal stem cells and samples untreated (negative control). These results were confirmed by the analysis of dermal papilla nuclei analyzed using ImageJ software 1.53. The graph in Figure 3 shows a significant increase in the measured parameter indirectly related to the number of cells in samples treated with product containing exosomes from colostrum bovine (45.886 ± 0.04) compared with product containing exosomes from mesenchymal stem cells (22.253 ± 0.03) and untreated cells (negative control) (18.112 ± 0.02).

**Figure 3.**
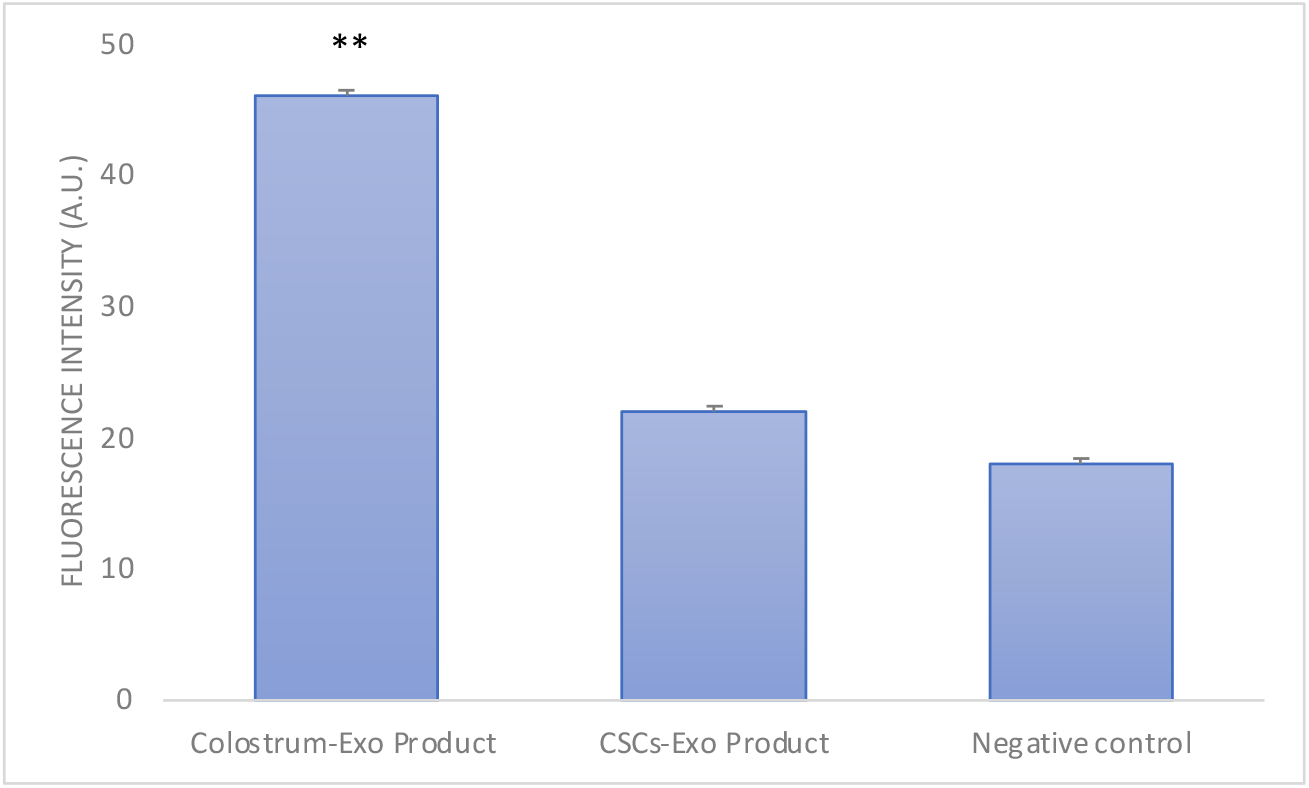
Analysis of the fluorescence intensity of the hair bulb via ImageJ. Comparation between two products contains exosomes. Statistically significantly different ** (p < 0.001).

The positive results observed for product containing exosomes from bovine colostrum compared to those containing exosomes from stem cells, could be related partly to the action of the exosomes and partly to that of the free factors present in product. The exosomes, in fact, through encapsulation into their structure, protect bioactive molecules from enzymatic and nonenzymatic degradation systems [19,20]. Exosomes beside have an important role in cell communication because they are recognized by target cells fusing with their membranes and releasing their contents [21]. But the effectiveness of this product is also due to the high concentration of growth factors and cytokines free which, by stimulating the anagen phase of the hair cycle and reducing the telogen phase of the hair cycle, help to restart the entire hair cycle [22,23].

## 4. Conclusion

Our study demonstrated a marked expansion of the germinative area in bulbs treated with the product containing colostrum derived exosomes compared to product containing stem cells derived exosomes, probably induced by activation of the Wnt/β-catenin pathway [11, 24, 25]. Exosomes that derived from bovine colostrum in fact can protect the hair follicles and stimulate and increase their growth. Therefore, this product could certainly be capable to combat hair loss.

## Author Contributions

Conceptualization, M.V.B.; methodology, M.V.B.; software, G.F. and M.Z.; formal analysis, G.F. and M.Z.; investigation, G.F.; data curation, G.F. and M.Z.; writing original draft preparation, M.V.B.; supervision, M.V.B. All authors have read and agreed to the published version of the manuscript.

## Institutional Review Board Statement

This study was performed in line with the principles of the Declaration of Helsinki and does not require approval from the Ethics Committee of University of Catania.

## Informed Consent Statement

Informed consent has been obtained from the volunteers to publish this paper.

## Data Availability Statement

Original data are available on request.

## Acknowledgments

G.F. thanks the Ph.D. program FSE Notice 1/2021.

